# Experimental recommendations for estimating lower extremity loading based on joint and activity

**DOI:** 10.1101/2021.05.14.444184

**Authors:** Todd J. Hullfish, John F. Drazan, Josh R. Baxter

**Author notes:** **CORRESPONDING AUTHOR** Josh R. Baxter, PhD. **AUTHORS’ CONTRIBUTIONS** TH and JB designed the experiment, TH performed the experiment, TH analyzed the data, all authors critically discussed the findings, TH and JB drafted the manuscript, all authors revised the intellectual content of the manuscript; all authors approved the final version of the manuscript.

## Abstract

Researchers often estimate joint loading using musculoskeletal models to solve the inverse dynamics problem. This approach is powerful because it can be done non-invasively, however, it relies on assumptions and physical measurements that are prone to measurement error. The purpose of this study was to determine the impact of these errors – specifically, segment mass and shear ground reaction force - have on analyzing joint loads during activities of daily living. We preformed traditional marker-based motion capture analysis on 8 healthy adults while they completed a battery of exercises on 6 degree of freedom force plates. We then scaled the mass of each segment as well as the shear component of the ground reaction force in 5% increments between 0 and 200% and iteratively performed inverse dynamics calculations, resulting in 1,681 mass-shear combinations per activity. We compared the peak joint moments of the ankle, knee, and hip at each mass-shear combination to the 100% mass and 100% shear combination to determine the percent error. We found that the ankle was most resistant to changes in both mass and shear and the knee was resistant to changes in mass while the hip was sensitive to changes in both mass and shear. These results can help guide researchers who are pursuing lower-cost or more convenient data collection setups.

## INTRODUCTION

Estimating joint loading during human movement is a cornerstone of biomechanics research. Traditionally, joint loads are estimated using musculoskeletal models to solve the inverse dynamics problem. Relying on Newton’s second law of motion, we sum the external forces acting on a body segment and set that equal to the body segment dynamics (Winter, 2009). This approach is powerful because it allows researchers to estimate the reaction loads at each joint that are impossible to physically measure without invasive surgeries (Bergmann et al., 2014, 2001). However, this approach relies on assumptions and physical measurements that are difficult to quantify and prone to measurement error.

Quantifying the mass of each body segment is a necessary step in solving the inverse dynamics problem. Pioneering work by Zatsiorsky and Seluyanov (1983) – which was later advanced by De Leva (1996) – developed mathematical equations to characterize the inertial properties of each body segment. Efforts continue to develop new methods to improve model assumptions and reduce measurement errors (El Habachi et al., 2015; Muller et al., 2017; Rao et al., 2006; Reinbolt et al., 2007). Advanced imaging is leveraged to develop subject specific musculoskeletal models (Davidson et al., 2008; Sheets et al., 2010), and techniques like the residual reduction algorithm reconcile dynamic inconsistencies between mass properties and experimental measurements (Delp et al., 2007; Langenderfer et al., 2008). However, these methods that improve inverse dynamics fidelity come with increased burdens that make wide scale measurements in prospective studies and fieldwork impractical due to added expense and complexity.

While accurately quantifying mass properties receives a great deal of attention in both the literature and classroom, the impact of the accuracy of these properties on analyzing joint loads during different activities of daily living is less clear. Therefore, the purpose of this study was to evaluate how the accuracy of joint load estimates derived from inverse dynamic calculations are impacted by errors in segment mass properties. To further explore the impact of experimental measurements on joint load estimates, we also tested the sensitivity to shear ground reaction force errors. We collected ‘gold-standard’ motion capture data on a group of healthy young adults across a variety of activities using a marker-based motion capture system and embedded 6 degree of freedom force plates and performed inverse dynamics to establish a ‘ground truth’ range for joint load estimates. We then systematically introduced error by manipulating the mass properties of the musculoskeletal model and the magnitude of the shear component of the externally applied loads. From this, we compared the peak joint load estimates from each error condition with the ‘ground truth’ across the joints in the lower extremities. We hypothesized that changing the mass properties and shear ground reaction forces would differentially impact estimated joint loading, with the smallest effects on the ankle and the greatest effects on the hip.

## METHODS

### Study Design

We recruited 8 healthy adults (6 males, 2 females; 30 ± 4 years; BMI, 24.1 ± 3.2 kg / m2) who provided written informed consent that approved by the University of Pennsylvania IRB. All procedures were performed in accordance with the relevant guidelines and regulations. We excluded participants if they had a history of lower extremity injury.

Participants wore standardized lab clothing (running shorts and tank top) and running shoes (Air Pegasus, Nike, Beaverton, OR), and we secured retro-reflective markers (9.5 mm, B&L Engineering, Santa Ana, CA) to the pelvis and lower extremities using skin-safe tape that we have described in a previous report (Slater et al., 2018). Briefly, we placed markers over anatomic landmarks of the pelvis: anterior and posterior superior iliac spines; legs: medial and lateral knee condyles and ankle malleoli; and feet: calcaneus, first and fifth metatarsal heads, and the great toe that were placed on the shoes. We also placed additional tracking markers on the pelvis and lower extremities: two markers on the sacrum, one marker on the thigh, and two markers on the shank. Once they were clothed and outfitted with the appropriate markers, we acquired a static trial with the participants standing in the anatomic position (Seth et al., 2018).

### Data Collection

Participants completed a battery of exercises that are clinically relevant for treating Achilles tendon injuries, as described in a previous paper from our group (Baxter et al., 2021). From this battery, we selected a subset of exercises that would provide a wide range of lower extremity loading for our later analysis. These exercises were single leg heel raises, forward single leg hopping, single leg drop jumps, squats, lunges, as well as walking and running at self-selected speeds. Participants completed 5 repetitions of the jumps, hops, squats, and lunges and 10 repetitions of the heel raises as well as enough walking and running trials to collect 10 foot-strikes each. To prevent fatigue, we provided participants 2-5 minute rest periods between each exercise.

While participants completed each exercise, we acquired marker trajectories using a 12-camera motion capture system (Eagle Series, Motion Analysis Corporation, Rohnert Park, CA) sampling at 100 Hz and ground reaction forces using 3-embedded force plates (BP600900, AMTI, Watertown, MA) sampling at 1,000 Hz. We later post-processed the motion capture data to prepare it for Inverse Dynamic analysis by confirming marker labeling, filling small gaps using cubic spline interpolation, and filtering marker trajectories using a low-pass Butterworth filter with a 6 Hz cutoff. We corrected for errors in the ground reaction forces using an established force plate calibration procedure (Collins et al., 2009).

### Data Analysis

We used a constrained kinematic model to calculate lower extremity kinematics and kinetics (Seth et al., 2018). First, we scaled a generic musculoskeletal model (gait2392) using each participant’s bodyweight and markers placed over anatomic landmarks. Next, we moved the scaled model into the anatomic position by fitting the experimental data collected during the static trial using best practices (Hicks et al., 2015). The markers placed on the anterior superior iliac spines, condyles and malleoli, calcaneus, 1st and 5th metatarsal heads, and toe markers were all given equal weighting. Similarly, the hip, knee, ankle, and toe joints were all weighted towards neutral sagittal alignments, which we visually confirmed during the static trial. We then confirmed the scaled models by superimposing the experimental marker positions over the model.

To test the effects of segment mass and shear ground reaction force errors, we iteratively performed inverse dynamics across a wide range of conditions. We first performed inverse kinematics and inverse dynamics to generate what we considered the ‘ground truth’ sagittal joint moments of the right leg at the hip, knee, and ankle during each activity. We then iteratively modified the segment mass properties and shear ground reaction forces to test the effects on inverse dynamics. We decided to scale the segment masses and shear ground reaction forces by 0 to 200% in 5% increments to establish the implications of a range of experimental conditions. For example, scaling each segment mass by 0% effectively eliminates the dynamics and represents a static solution. Scaling the shear ground reaction forces by 0% tests the potential fidelity of using low-cost force plates that only measure the vertical ground reaction force. We further tested the interaction between segment mass properties and shear ground reaction forces on joint load estimates by running inverse dynamics using each mass-shear combination. In total, we performed 1,681 mass-shear combinations for each movement trial. We calculated the percent error in peak joint moments between the ‘ground truth’ condition and each of the mass-shear combinations as our primary outcome measure. Using these percent errors, we generated heat maps to visualize the interaction between segment mass and shear ground reaction forces on joint moments.

To better understand the real-world implications of these simulations we compared four mass-shear combinations that represent potential data collection setups: 1) traditional motion capture with marker-based kinematics and embedded 6 degree of freedom force plates (100% mass, 100% shear force); 2) marker-based kinematics and vertical component force plates (100% mass, 0% shear force); 3) marker-less pose estimation kinematics and 6 degree of freedom force plates (0% mass, 100% shear force); and 4) marker-less pose estimation kinematics and vertical component force plates (0% mass, 0% shear force). By assessing these peak joint moment errors at the hip, knee, and ankle across a range of activities of daily living, we established guidelines for when body segment masses or ground reaction force data must be carefully attained and when simplified data analyses and experimental setups are justified. Here we based our recommendations on ‘real-world’ experimental setups that preserved joint load fidelity within 10% of the ‘gold-standard’ of 100% mass and 100% shear ground reaction force calculations.

## RESULTS

Joint moment errors tended to increase in more proximal joints based on shear ground reaction force errors more than segment mass errors (Figure 1). The ankle was the least sensitive to errors in both segment mass and shear ground reaction force, experiencing an average error in peak plantar flexion moment of 10% across all activities. Walking and lunging had the greatest peak plantar flexion moment errors of 18 and 28%, respectively, while running and heel raising had small peak plantar flexor moment errors of around 2%. The hip was the most sensitive, with an average peak hip extension moment error of 77% across all activities. Walking and single leg hopping generated the largest errors in peak hip extension moment of 117 and 172%, respectively, while drop jumping resulted in the smallest peak hip extension moment error of 18%. The knee was less sensitive than the hip but still experienced large errors with an average of 56%. Like the hip, walking and single leg hopping resulted in the largest errors in peak knee extension moment of 148 and 86%, respectively, while squatting resulted in the smallest error of peak knee extension moment of 7%. The knee was more sensitive to errors in shear ground reaction forces while the hip was more sensitive to errors in segment mass. On average, neither shear ground reaction forces nor mass impacted peak plantar flexion moments more than 10%. However, walking and lunging were more sensitive to errors in shear ground reaction forces.

**Figure 1.**
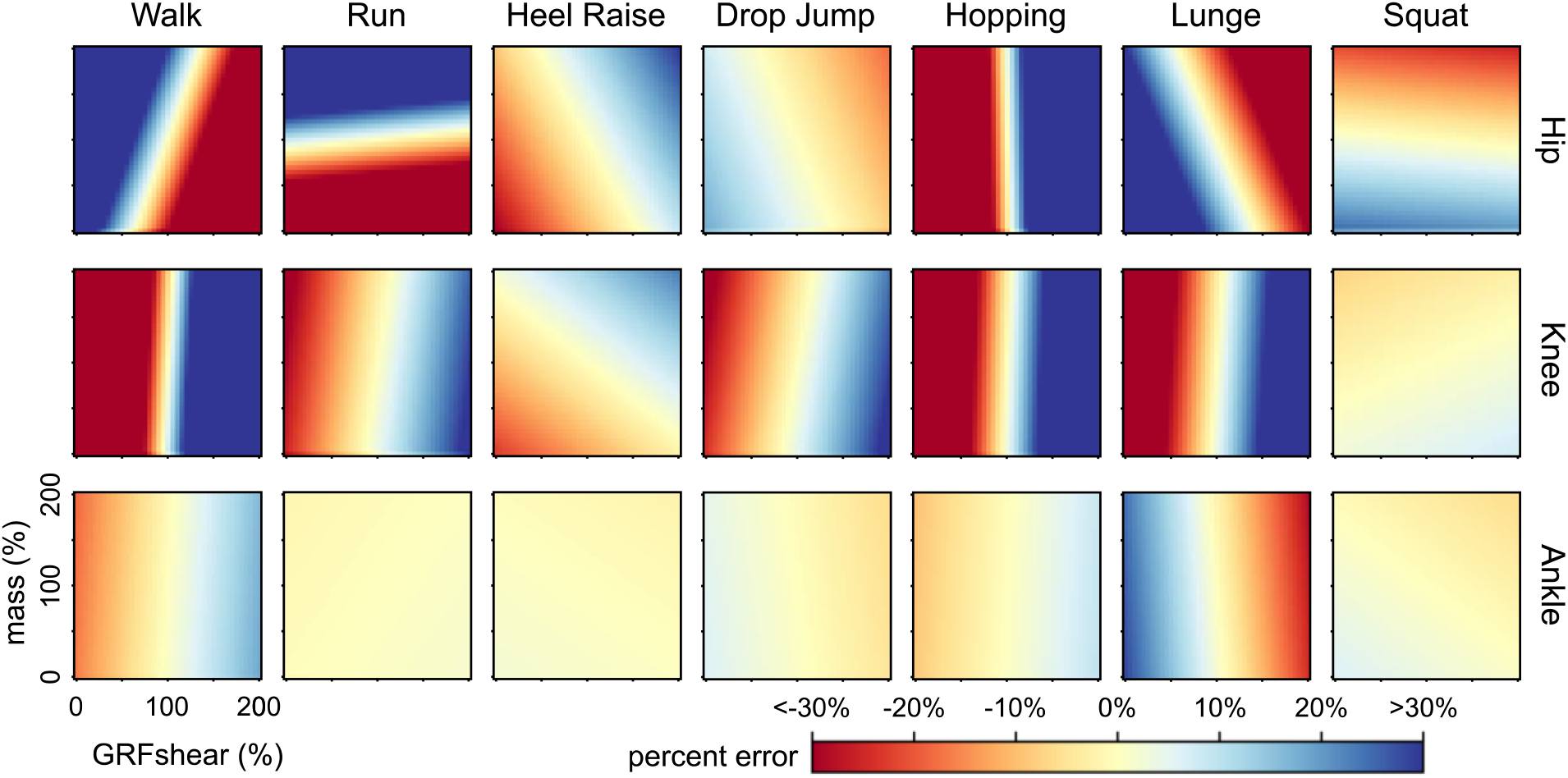
Errors in peak joint moment for each mass-shear combination visualized as heat maps for the hip, knee, and ankle for each of the analyzed motions. Each pixel of a given heat map represents the percent error of peak joint sagittal moment between the mass-shear combination and the ground truth combination (100% mass – 100% GRFshear). Pale yellow represents 0% error, deep red represents <-30% error, and deep blue represents >30% error.

We established experimental setup recommendations specific to activity type and joint of scientific interest (Figure 2). We found that accurate segment masses were unnecessary to accurately quantify ankle joint moments in all activities. Activities like heel raises and vertical hopping were not sensitive to inaccurate shear ground reaction forces (Figure 2). However, activities that generated considerable amounts of shear ground reaction forces like walking and lunging caused ankle joint loading errors in excess of 28%. Knee joint loading was sensitive to shear ground reaction forces but not segment mass. For example, there was less than 10% error in 6 out of 7 activities when shear ground reaction forces were accurate compared to only 2 out of 7 when segment masses were accurate. Hip joint loading was sensitive to even small errors in both ground reaction shear and segment mass and there were only 5 out 21 conditions where the error was less than 10% (Figures 1 and 2).

**Figure 2.**
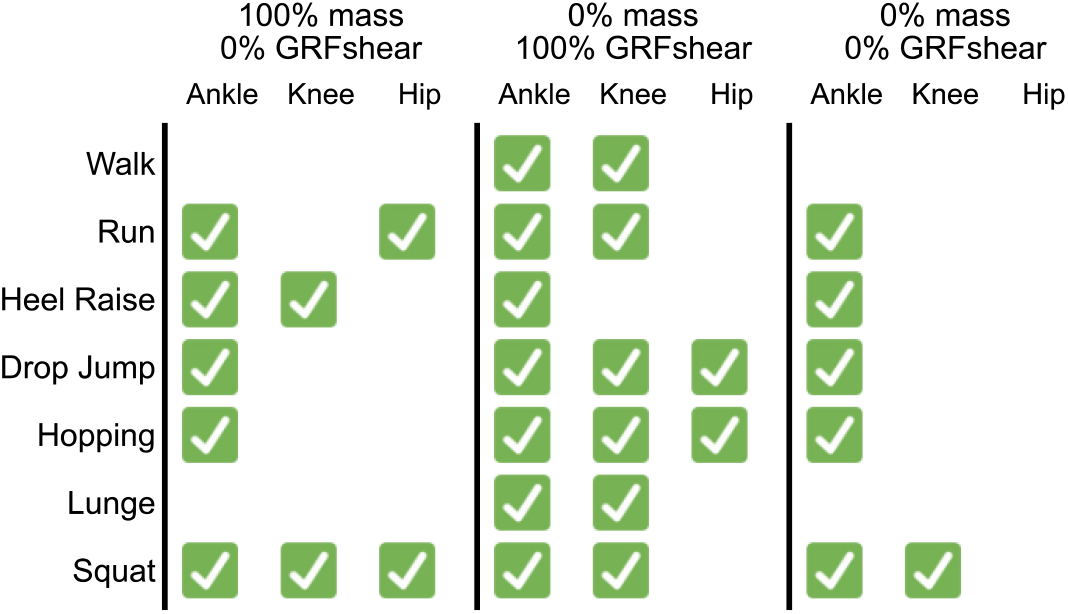
Experimental recommendations to achieve desired joint loading fidelity. Scenarios where there was less than 10% error compared to ‘ground truth’ calculations from inverse dynamics(100% mass and 100% GRFshear) are demarked with a check mark. 0% mass conditions represent experimental techniques that quantify body ‘pose’ and do not quantify segmental dynamics. 0% GRFshear represent experimental conditions that may utilize low-cost force plates that only measure the vertical reaction force.

## DISCUSSION

The purpose of this study was to determine the effects of errors in segment mass and ground reaction shear forces on inverse dynamic calculations of peak hip, knee, and ankle moments. As we hypothesized, the ankle was least sensitive to these errors while the hip was the most sensitive. Accurate ground reaction force measurements appear to be more important than accurate measurements of segment mass in most situations for calculating peak joint moments through inverse dynamics. These findings agree with our intuition that changing the direction of the ground reaction force will have greater impacts on joint loading than changes in the dynamic forces. For example, the ankle joint experiences ground reaction forces in excess of 1.2 times bodyweight during the stance phase of walking which far exceed the dynamic forces of the foot accelerating over the ground. These results provide quantitative support for employing lower cost experimental setups without sacrificing biomechanical fidelity in some instances.

Our results are promising for researchers who are exploring more practical experimental setups while preserving high-fidelity analysis (Figure 2). For example, ankle and knee loads can be considered quasi-static while the foot is contacting the ground, which opens up new opportunities for leveraging pose-estimation techniques to analyze relevant instances of a movement rather than relying on high-speed motion capture to fully characterize movement dynamics. Experimental setups can be simplified further when analyzing movements that mostly change the height of the center of mass. Movements like vertical hopping and heel-raises are resistant to shear ground reaction force errors (Figure 1). In these special scenarios, quasi-static analyses using low-cost vertical-component force plates provide similar levels of fidelity as traditional laboratory setups.

Understanding how measurement errors impact inverse dynamics calculations is critical as the field continues to explore research questions that are best studied outside of traditional biomechanics laboratory. Estimating joint loading using low-cost motion capture techniques or wearable devices has emerged as promising tools to study patients in more natural settings, both in the clinic and in the real world (Hullfish et al., 2020; Matijevich et al., 2020; Renner et al., 2019). These approaches compare favorably to those made using gold-standard techniques across a wide range of clinically relevant activities (Drazan et al., 2021; Hullfish and Baxter, 2020; Martin et al., 2018). Most of these studies have focused on either the knee or the ankle and our results seem to support the validity of these paradigms for producing high quality data.

However, our results should be cautiously applied to each research question. Leveraging gold-standard laboratory equipment to maximize measurement fidelity is prudent for applications that impact clinical movement analyses used to inform patient care like surgical planning (Arnold and Delp, 2005) or assistive devices (Guan et al., 2016). In these scenarios, where the measurements inform subject specific treatment or assistive devices, the increased costs associated with lab-based techniques are clearly justified. Additionally, research questions focused on the hip joint should treat most activities as dynamic and measure ground reaction forces using 6 degree of freedom force plates.

This study had limitations that are important to consider when putting the results in context. We assumed that our musculoskeletal models represented accurate segment masses for each subject. While we didn’t employ advanced scaling techniques (Killen et al., 2021; Valente et al., 2017), our sensitivity analysis tested extremes that far exceeded any plausible differences between the segment masses defined in the musculoskeletal model and subject-specific segment masses. We focused our analyses on instances when the foot is contacting the ground because anytime the foot is off the ground, joint loading errors will be directly proportionate to segment mass errors. We also focused on sagittal joint loads to establish the efficacy of using existing pose-estimation techniques to quantify joint loading and because sagittal movements represent the greatest amount of body accelerations and shear ground reaction forces. We also note that knee and hip errors during activities that are predominantly plantar flexion movements – in this study, heel raise and hopping – appear large because ‘ground truth’ peak loads are small. For example, peak knee loading during a heel raise is only 0.18 Nm/kg compared to 1.99 Nm/kg during the stance phase of running.

In summary, we performed a sensitivity analysis to test the effects of changing body segment masses and shear ground reaction forces on lower extremity joint moment calculations across a range of clinically relevant activities. In general, we found that ankle and knee joint loads were resistant to changes in mass properties – effectively confirming that quasi-static solutions can replace inverse dynamics for many applications. Conversely, hip joint loading is sensitive to both segment mass and shear ground reaction forces, highlighting the need for high-fidelity measurements. We used these analyses to develop guidance for utilizing less costly – in terms of financial and logistical – experimental setups based on the joint and activity of research interest. We hope that this guidance benefits the biomechanics community as we continue to develop innovative new techniques to quantify joint loading outside of traditional biomechanics laboratories.

## ACKNOWLEDGEMENTS

This work was supported by the PennPORT IRACDA Fellowship Program (NIH grant# K12GM081259) and NIH/NIAMS K01AR075877.

